## Supplementary Figure 1-4 for "Fast imaging of millimeter-scale areas with beam deflection transmission electron microscopy"

Supplementary Fig. 1

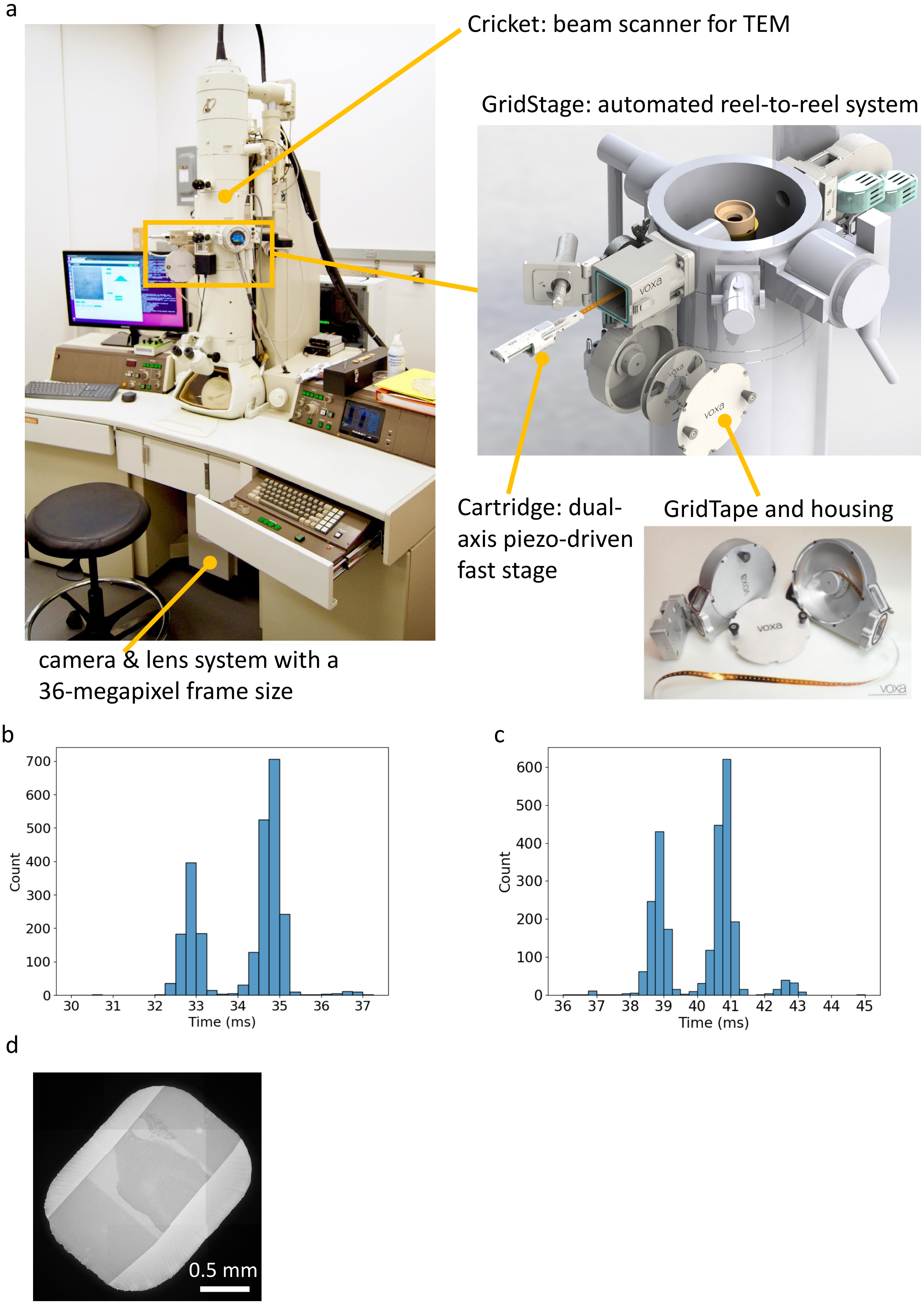

Supplementary Fig. 2

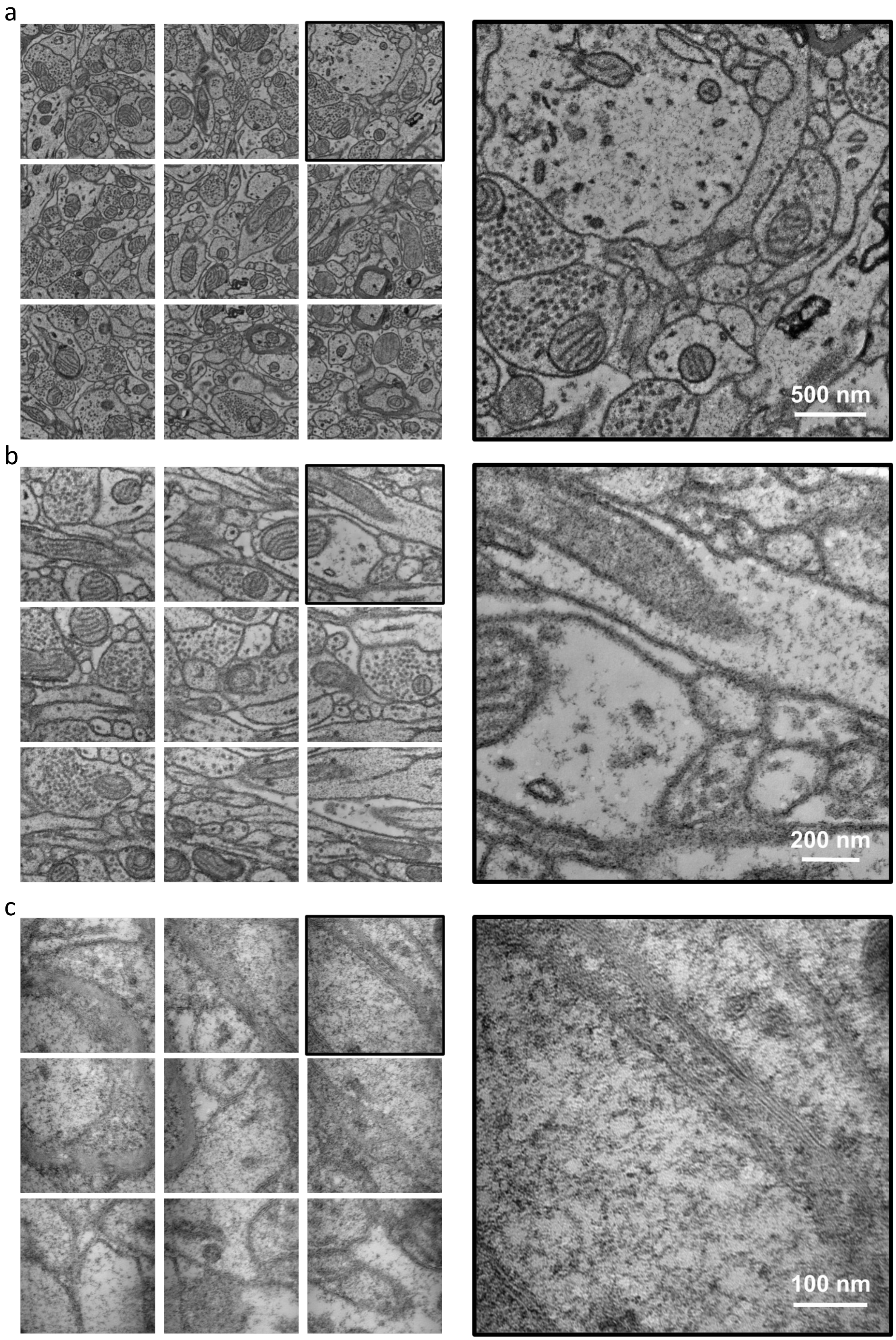

Supplementary Fig. 3

a

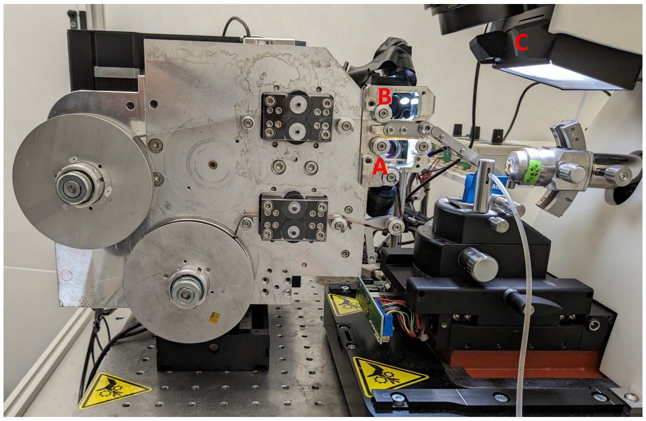


b

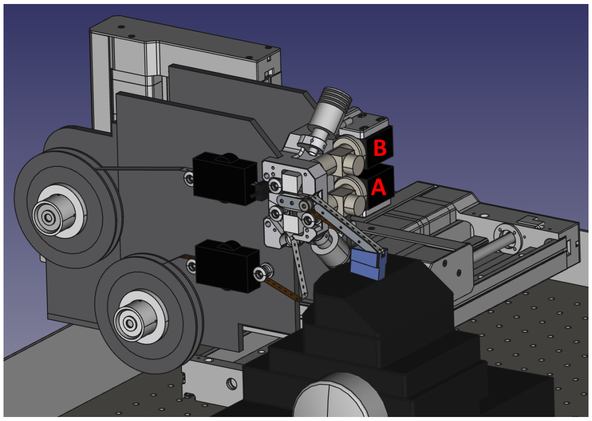


c

Tape control | Water level control | Tape stage control

A. before sectioning | B. after sectioning | C. during sectioning

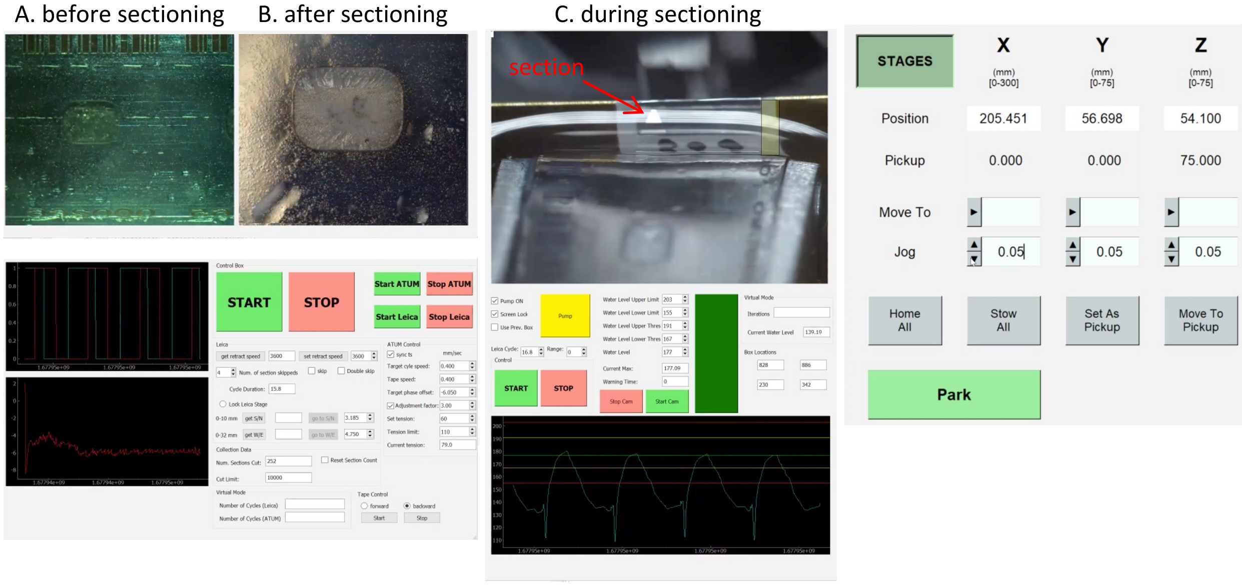

Supplementary Fig. 4

a

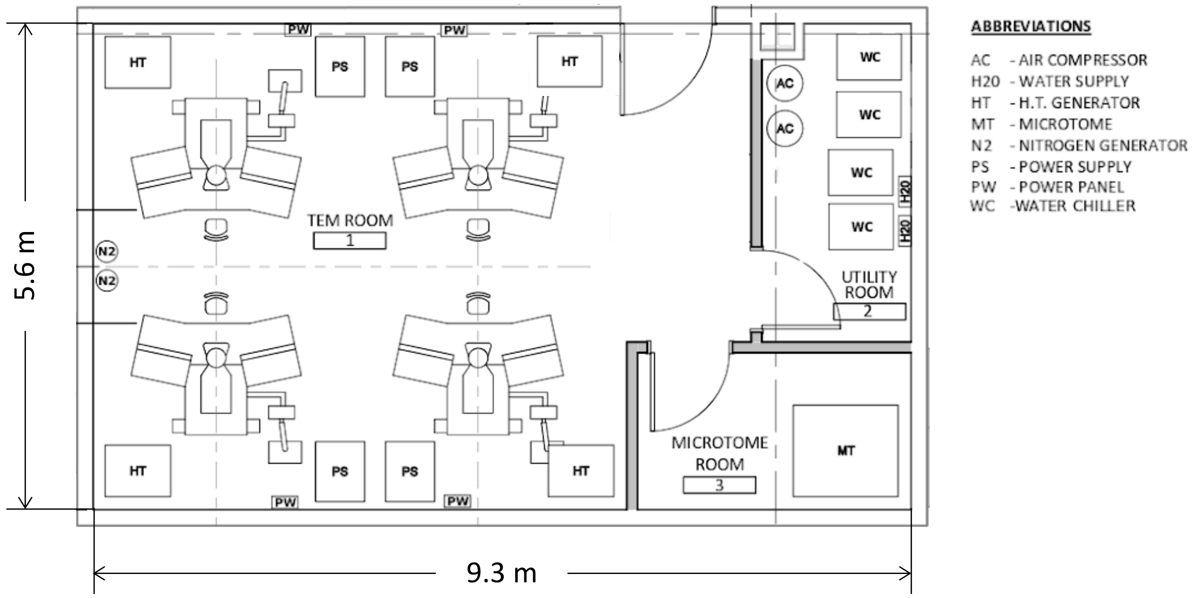


b

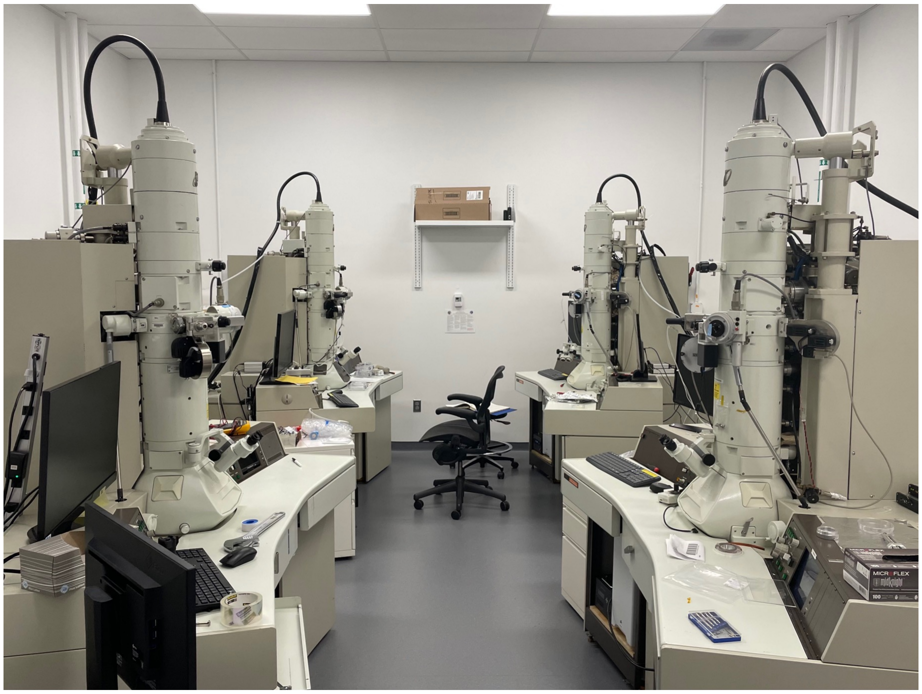
